# Quantifying Waddington’s epigenetic landscape: a comparison of single-cell potency measures

**DOI:** 10.1101/257220

**Authors:** Jifan Shi, Andrew E. Teschendorff, Weiyan Chen, Luonan Chen, Tiejun Li

## Abstract

Over 60 years ago Waddington proposed an epigenetic landscape model of cellular differentiation, whereby cell-fate transitions are modelled as canalization events, with stable cell states occupying the basins or attractor states^1, 2^. A key ingredient of this landscape is the energy potential, or height^3^, which correlates with cell-potency. To date, very few explicit biophysical models for estimating single-cell potency have been proposed. Using 9 independent experiments, encompassing over 6,600 high-quality single-cell RNA-Seq profiles, we here demonstrate that single-cell potency can be approximated as the graph entropy of a Markov Chain process on a model signaling network. Our analysis highlights that other proposed single-cell potency measures are not robust, whilst also revealing that integration with orthogonal systems-level information improves potency estimates. Thus, this study provides a foundation for an improved systems-level understanding of single-cell potency, which may have profound implications for the discovery of novel stem-and progenitor cell phenotypes.

---

According to Waddington’s epigenetic landscape model of cellular differentiation, human embryonic stem cells (hESCs) occupy the highest attractor state within this landscape, owing to their pluripotency, with terminally differentiated cells occupying the lowest lying basins. Quantifying the energy potential, or potency, of single-cells is a task of critical importance, as this is necessary to allow explicit construction of Waddington landscapes^2,4^, but also to provide an unbiased means of identifying novel stem or progenitor cell phenotypes.

Given that mRNA expression levels of genes inform on the activity of transcription factors and signaling pathways which control cell potency^9^, it is reasonable to assume that potency is encoded by the genome-wide transcriptomic profile of the cell^5–8^. Following this rationale, a number of explicit potency models based on biological principles have been proposed^6–8^. For instance, SLICE^7^ defines potency in terms of a Shannon entropy over gene expression derived gene-ontology activity estimates, which in effect measures the relative activation profile of different biological processes in a cell. StemID^6^ computes a genome-wide transcriptomic entropy which measures how uniformly expressed the genes are. SCENT^8^ models potency in terms of the entropy-rate of a diffusion process^10^ on a signaling network, which measures how efficiently signaling can diffuse over the whole network. Here, we introduce a novel single-cell potency measure, called Markov-Chain Entropy (MCE) (Fig.1A, Online Methods). Like SCENT, MCE integrates the RNA-Seq profile of the single-cell with a model (protein-protein) interaction network, but differs from SCENT in that it models potency as the graph entropy of an optimized Markov Chain process whose invariant measure is the observed steady-state gene expression profile (Fig.1A, Online Methods).

**Figure 1.**
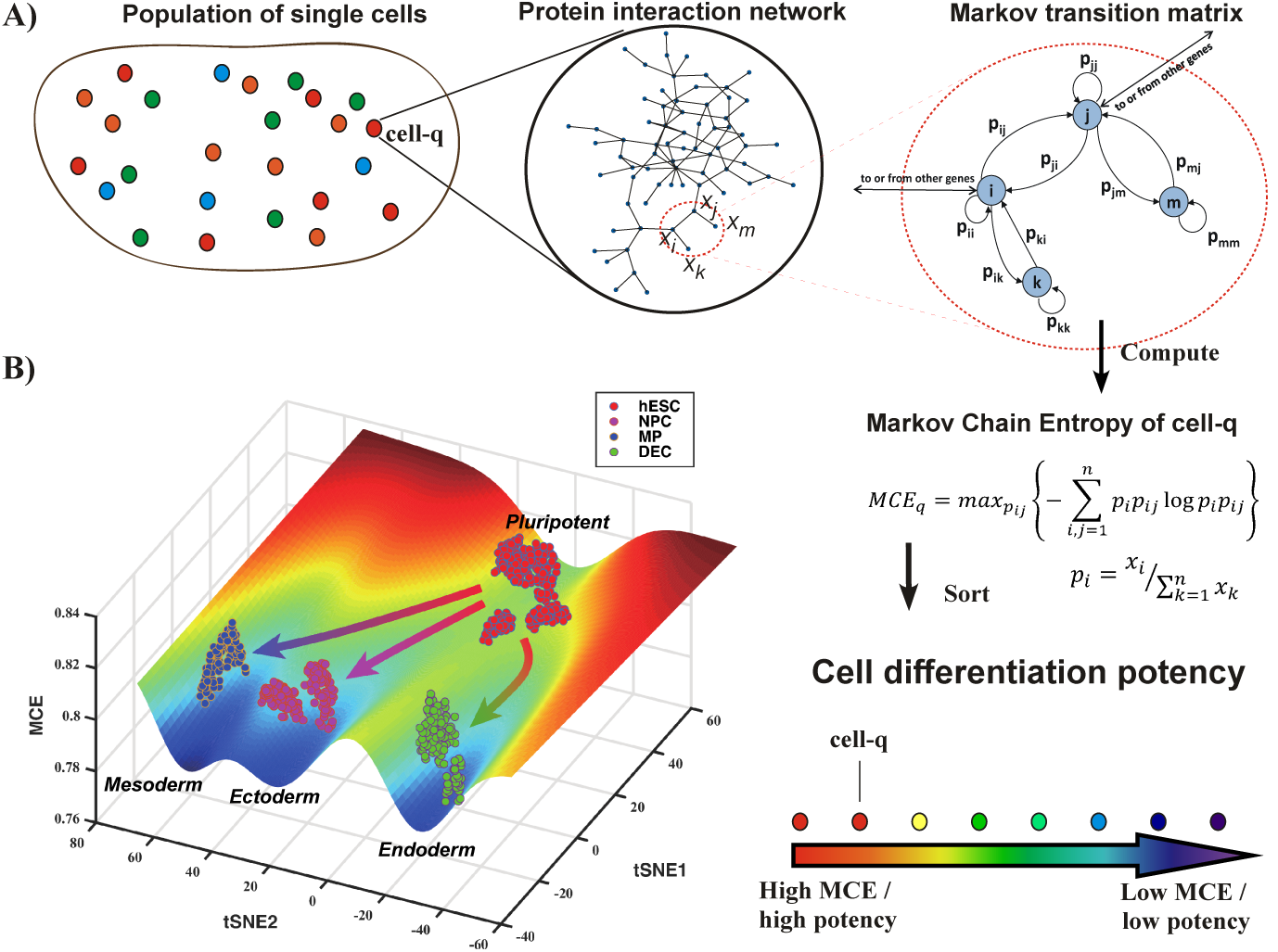
Markov-Chain Entropy for quantifying single-cell potency in Waddington’s landscape. **A)** The key assumption underlying the estimation of Markov-Chain-Entropy (MCE) is that the measured gene expression profile ***x*** = (*x*_1_, …, *x*_n_) of a cell q is the end result of an unknown Markovian signaling process in the cell, which however we can try to estimate by using a network model of signaling interactions. This network imposes constraints on the space of all possible Markov transition matrices, and MCE is estimated from the one maximizing the Shannon entropy of the Markov process. Having estimated MCE for each single cell, we can then sort them according to their MCE, which we posit is a proxy for cell potency. **B**) An example of a Waddington epigenetic landscape for the Chu1 et al. data set (**Table S1 in SI**), depicting 4 main attractor states, representing hESCs (pluripotent), neural progenitor cells (NPCs, ectoderm, multipotent), mesoderm progenitors (MPs, mesoderm, multipotent) and definite endoderm progenitor cells (DECs, endoderm, multipotent). The MCE values of over 1000 single cells representing these 4 cell-types are represented above the attractor states.

To assess the ability of MCE to measure single-cell potency and to benchmark it to previously proposed measures, we devised an objective evaluation framework: we reasoned that an objective performance criterion is the ability to discriminate single cells that should differ significantly in terms of potency, for instance, when comparing pluripotent to non-pluripotent cell populations, or between cells at the start and end points of time-course differentiation experiments. Thus, building on recent advances in single-cell RNA-Seq (scRNA-Seq) technology^3,11–13^, we assembled a collection of 9 scRNA-Seq experiments, encompassing a total of approximately 6700 high-quality scRNA-Seq profiles, from which we devised 11 independent objective comparisons between cell-types that ought to differ in terms of potency (Online Methods, **Table S1 in SI**). To each of these datasets, we applied the three previously proposed single-cell potency models (StemID^6^, SLICE^7^, SCENT^8^), as well as MCE.

We first considered a time-course differentiation experiment in which hESCs were induced to differentiate into cortical neural progenitors and finally into neurons over a 54-day period, with scRNA-Seq measurements taken at 6 time points, encompassing a total of 2684 single-cells^14^. While MCE and SCENT showed the expected gradual decrease of potency with differentiation time point, StemID exhibited a less significant pattern, whilst SLICE did not exhibit a decrease (Fig.2A). Comparing hESCs to cortical neural progenitors and then to differentiated neurons separately, revealed that the potency measures in SCENT and MCE could discriminate cells with almost perfect accuracy (Fig.2A-B, **Table S2**). As expected the difference in potency between progenitors and neurons was less strong, but still significantly stronger than the differences predicted by StemID or SLICE (Fig.2A-B, **Table S2**). As assessed over the 11 independent comparisons (**Table S1 in SI**), we observed that the MCE and SCENT potency measures outperformed the other two (SLICE and StemID), also when adjusted for cell-cycle phase (Fig.2B-C, **Table S2-S3 in SI**, Online Methods). In 9/11 comparisons, MCE and SCENT were better than StemID, and in all 11 better than SLICE. The better performance of MCE and SCENT was statistically significant under a paired Wilcoxon test over the 11 comparisons (Fig.2B-C).

**Figure 2.**
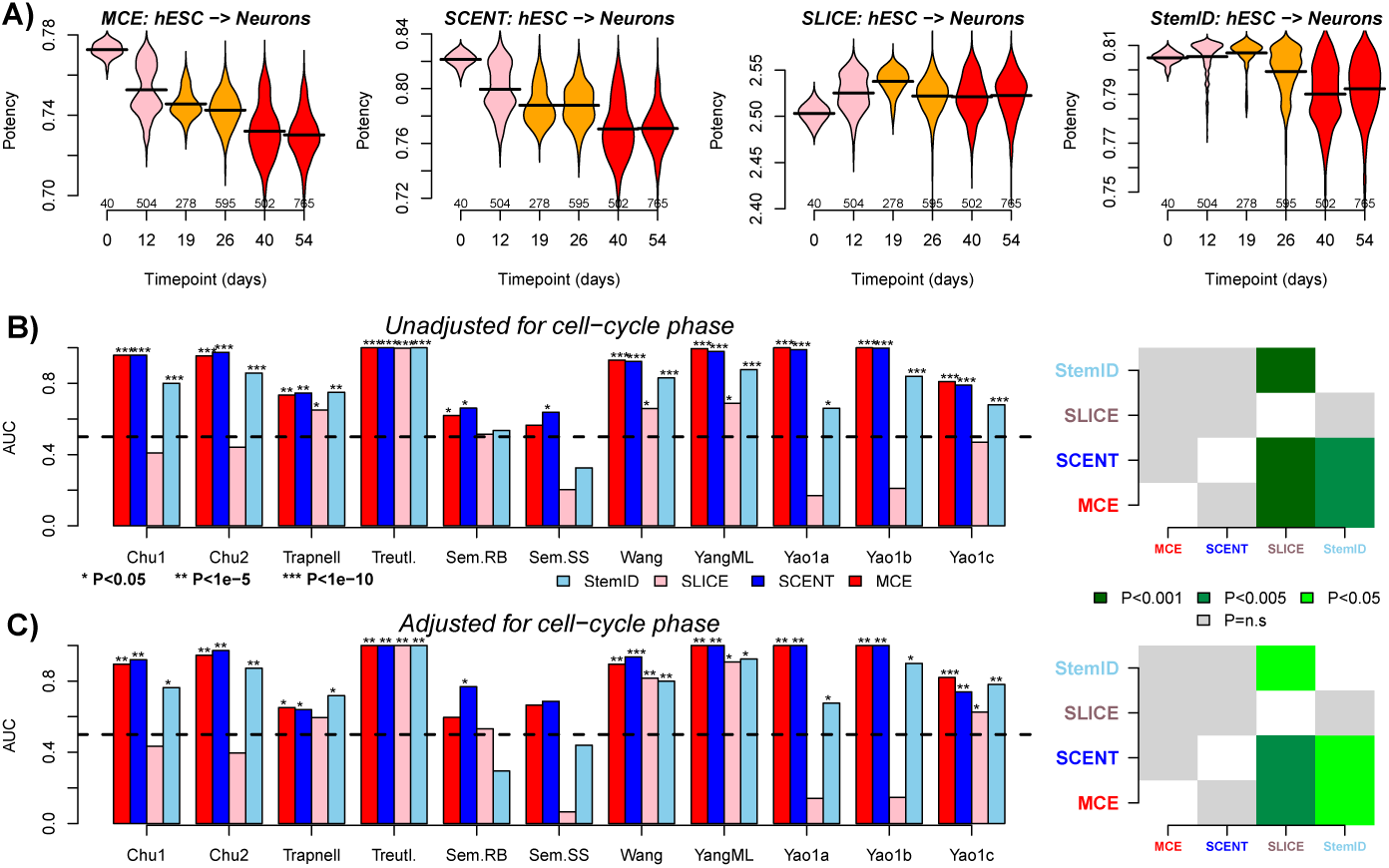
Comparison of discrimination accuracy of single-cell potency measures across scRNA-Seq datasets. **A**) For a scRNA-Seq dataset profiling over 2600 cells in a time course differentiation of hESCs (day-0) to cortical neural progenitors (day 26) and finally to terminally differentiated neurons (day-54), we show corresponding violin plots of the predicted potencies at each measured timepoint, and for the 4 different potency measures (SLICE, StemID, MCE and SCENT), as indicated. The number of cells at each timepoint is given below violins. **B) Left panel:** Barplots compare the Area Under the Curves (AUCs) of the four different single-cell potency measures across 11 independent comparisons drawn from 9 independent scRNA-Seq experiments. The AUCs reflect the measure’s discrimination accuracy of single cells that ought to differ in terms of differentiation potential, and can be derived from the statistic of a Wilcoxon rank sum test. See Table S1 for definitions of cell-types being compared. P-values are derived from a one-tailed Wilcoxon rank sum test comparing cell-types. **Right panel:** Heatmap of meta-analysis P-values comparing single-cell potency measures to each other. P-values estimated from a paired one-tailed Wilcoxon rank sum test over the 11 independent comparisons. In the heatmaps, the P-value entry for column i and row j is for testing the alternative hypothesis that method in row j has higher AUC values than method in column i. **C**) As B), but now adjusting for cell-cycle phase, i.e. using only the cells with the lowest cycling scores in each comparison group.

Since potency is also a property of a cell population^8,15^, any robust measure of single-cell potency should, in principle, also work for bulk samples since a bulk RNA-Seq profile is effectively an average over single-cells^5^. We observed that only SCENT and MCE exhibited a robust performance on bulk-samples (**Table S4 in SI**), in line with the results obtained on scRNA-Seq data. Thus, based on the comparative analysis performed here, we conclude that SCENT and MCE currently provide the more robust measures of cell-potency, and therefore advocate their use for explicit, unbiased, construction of Waddington landscapes. As an example, we used MCE on the Chu1 et al. dataset (**Table S1 in SI**, Online Methods) in combination with t-stochastic neighborhood embedding (t-SNE) to construct a Waddington epigenetic landscape encompassing 4 attractor states, representing pluripotent hESCs and multipotent (i.e. non-pluripotent) progenitors from the 3 main germ layers: ectoderm, mesoderm and endoderm (Fig.1B). We also generated 3-dimensional representations for (i) an embryonic time-course differentiation experiment of mouse hepatoblasts into hepatocytes and cholangiocytes (Yang et al., **Table S1 in SI**), which correctly predicted separation after embryonic day-13 (Fig.3A), and (ii) for a 30-day time course differentiation experiment of neural progenitor cells into neurons (Wang et al., **Table S1 in SI**), where MCE correctly predicted a gradual decrease in potency Fig.3B).

**Figure 3.**
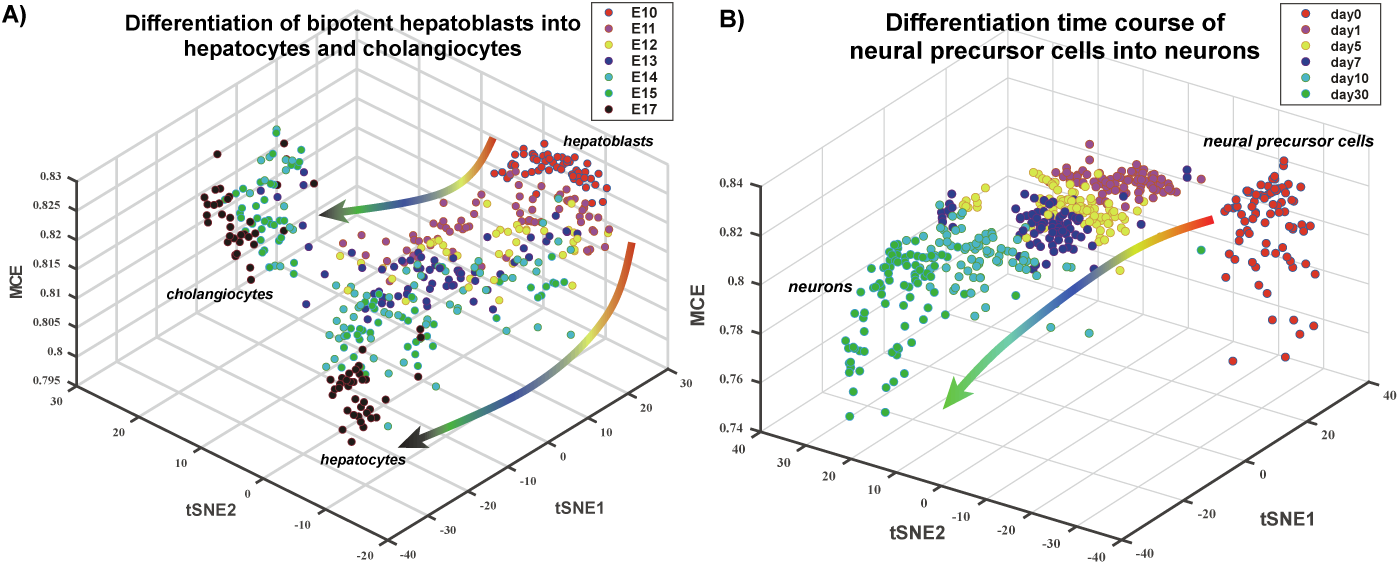
Three dimensional representation of single-cell differentiation trajectories according to Markov Chain Entropy. **A**) Differentiation landscape of hepatoblasts. Figure depicts a 3-dimensional representation of single-cells in an embryonic differentiation time course of mouse hepatoblasts into hepatocytes and cholangiocytes (Yang et al), as indicated. The MCE potency estimate labels the z-axis, whereas the x, y axes correspond to the t-stochastic neighborhood embedding (tSNE) components. Observe how the model predicts separation/differentiation into hepatocytes and cholangiocytes at embryonic day-13 (E13) in line with the results of Yang et al. **B**) Differentiation landscape of neural precursor cells into neurons. Figure depicts a 3-dimensional representation of single-cells in a differentiation time course of human neural precursor cells (day-0) into neurons (day-30) (Wang et al), as indicated. Observe how MCE correctly predicts a gradual decrease in potency with time of differentiation.

In order to understand the increased robustness of MCE and SCENT over StemID and SLICE, we note that both MCE and SCENT integrate the scRNA-Seq data with orthogonal connectivity information as derived from a protein-protein interaction network^8^ (Fig.1A, Online Methods). While such networks are mere caricatures of the complex spatially and temporally dependent signaling processes that take place in a cell, it is plausible that potency is encoded by specific robust features of such networks (e.g. the difference in connectivity between a hub and a low-degree node). As shown by us previously in the context of SCENT, its robustness as a potency measure derives from it being encoded by a subtle positive correlation between the RNA-Seq profile of a sample (be it a single cell or bulk) and the connectivity profile as defined in a PPI network^8^. Under this model, cells that express higher levels of highly connected genes (i.e. network “hubs”) will exhibit higher potency. We further demonstrated that such genes encode not only stemness genes but importantly also mRNA splicing factors and genes encoding mitochondrial ribosomal proteins^8^, consistent with studies demonstrating that mitochondrial activity and splicing rate influence stemness and differentiation^16–23^. Here we verified that the network is a critical feature underlying the robustness of MCE as a potency measure (Fig.4A-C, **Fig.S1 in SI**) and that this is partly driven by the correlation between transcritome and connectome (Fig.4D-E, **Fig.S1 in SI**). Thus, the increased robustness of MCE/SCENT owes to three separate properties: (i) the relative connectivity between hubs and low-degree nodes in a network is a robust feature of such networks, (ii) big changes in gene expression, specially at hubs, will have a larger influence on the MCE/SCENT estimates, and the larger changes in mRNA expression are more likely to be robust, and (iii) MCE/SCENT are computed genome-wide over a large number of genes, which renders them robust to the potentially large numbers of dropouts in scRNA-Seq data^13^.

**Figure 4.**
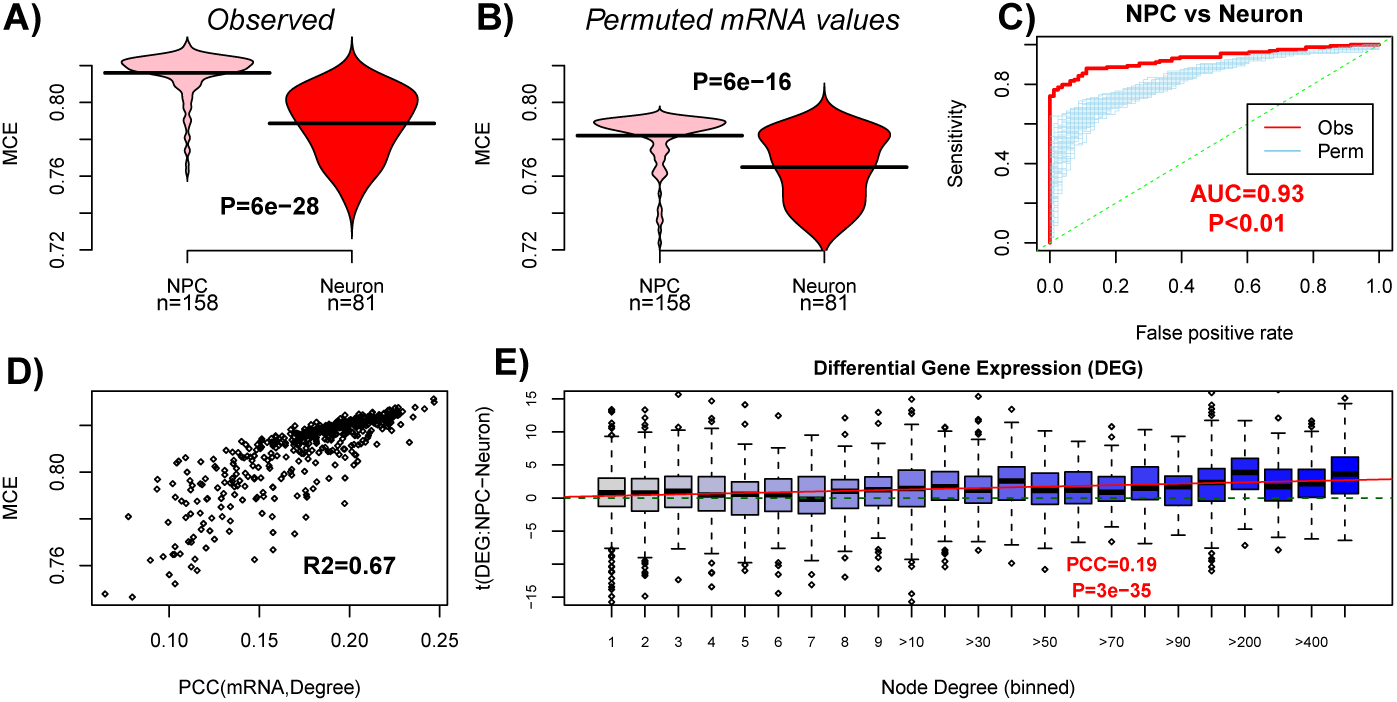
Integration with network information underpins the association of MCE with cell-potency. **A**) Violin plots of the observed MCE values for neural progenitor cells (NPCs) and terminally differentiated neurons (scRNA-Seq data from Wang et al). P-value derives from a one-tailed Wilcoxon rank sum test. **B**) As A), but now for a random permutation of the expression values over the network, leading to a reduced discrimination accuracy. **C**). Comparison of the Area Under the Curve (AUC) discrimination measure for the observed case (depicted in A)) and for 100 distinct permutations. P-value is an empirical one comparing the observed AUC to those from the 100 permutations. **D**) MCE (y-axis) is approximated reasonably well by the Pearson Correlation Coefficient (PCC) between the transcriptome and connectivity/degree profile of the network (x-axis). *R*^2^ value is given. **E**) Boxplots of t-statistics of differential expression between NPCs and neurons (y-axis) against node-degree (x-axis), with larger degrees binned into equal sized groups. Green dashed line denotes the line t=0. Red line is that of a linear regression. PCC is the Pearson Correlation between the t-statistics and node-degree. We also give the P-value for the linear regression.

We end by noting that the correct ordering of single-cell potency by MCE/SCENT was achieved for each cell individually, starting out from a purely model-driven approach, without using any prior biological information (e.g. known marker genes, sampling timepoint) and without using information from other cells (i.e. no feature selection was done, nor is clustering used to group cells). This is in stark contrast to most single-cell algorithms^24–32^, whose main aim is to infer cell-lineage trajectories from time-course differentiation experiments, and which all use either prior information (e.g. known marker gene expression, timepoint), or heuristics that lack strong biological justification, to assign cells to potency states. We therefore propose that future single-cell analysis algorithms should focus more on developing and testing improved in-silico models of cell-potency, as this is a key step towards a more accurate and unbiased reconstruction of Waddington landscapes^4^.

## Acknowledgements

This work was supported by NSFC (National Science Foundation of China) grants 31571359, 31401120, 91529303, 31771476, 11421101 and 91530322, by a Royal Society Newton Advanced Fellowship (NAF project: 522438, NAF award: 164914). This work was also supported by National Key R & D Program of China (2017YFA0505500).

## Author Contributions

Manuscript was conceived and written by TL, CL, AET and SJ. WC contributed to analyses.

## Online Methods

### Single cell and bulk RNA-Seq datasets

In total, we used single-cell RNA-Seq datasets (scRNA-Seq) derived from 9 independent studies. Two of these studies contained matched bulk RNA-Seq data. From all of these scRNA-Seq studies, we devised 11 independent analyses, comparing in each one cells of high potency to cells of low potency, assessing the different potency measures in their ability to discriminate these cell-types. In the case of bulk RNA-Seq data, we devised 5 independent comparisons from the two studies with bulk RNA-Seq data. Table S1 in SI gives a brief summary of the 11 independent comparisons performed on scRNA-Seq data and the 5 performed using bulk RNA-Seq data. A more detailed description of the datasets is listed below:

- *Chu et al. datasets 1-4:* These datasets were derived from Ref. 33. There are four experiments named Chu1, Chu2, Chu3 and Chu4. Chu1 dataset contains scRNA-Seq data for 1018 single cells, which is composed of 374 hESCs (human embryonic stem cells, 212 from H1 cell line and 162 from H9 cell line), 173 NPCs (neuronal progenitor cells, ectoderm derivatives), 138 DECs (definitive endoderm cells, endoderm derivatives), 105 ECs (endothelial cells, mesoderm progenitors/derivatives), 69 TBs (trophoblast-like cells, extraembryonic derivatives) and 159 HFFs (human foreskin fibroblasts). The hESCs are pluripotent (*n* = 374) and the others are non-pluripotent (*n* = 644). Chu2 dataset is a time course differentiation of single cells, in which hESCs were induced to differentiate into DECs, via a mesoendoderm intermediate. The time points cover before induction at *t* = 0h (*n* = 92) and after induction at *t* = 12h (*n* = 102), *t* = 24h (*n* = 66), *t* = 36h (*n* = 172), *t* = 72h (*n* = 138) and *t* = 96h (*n* = 188). There are 758 single cells in total. Chu3 dataset is from bulk samples. There are 19 bulk samples, which have 7 samples for hESCs, 2 for NPCs, 2 for TBs, 3 for HFFs, 3 for ECs (mesoderm progenitors) and 2 for DECs. Chu4 is the time-course experiment of 15 bulk samples, consisting of 3 samples for each of five time points (12h, 24h, 36h, 72h and 96h). Normalized data was downloaded from GEO under accession number GSE75748 (files: GSE75748-bulk-cell-type-ec.csv, GSE75748-sc-cell-type-ec.csv, GSE75748-bulk-time-course-ec.csv, GSE75748-sc-time-course-ec.csv).
- *Trapnell et al. dataset:* This scRNA-Seq dataset was derived from Ref. 24. It represents a time-course differentiation experiment of human myoblasts into human skeletal muscle cells. Human myoblasts were induced at *t* = 0h (*n* = 96). Samples were also taken at *t* = 24h (*n* = 96), *t* = 48h (*n* = 96) and *t* = 96h (*n* = 84) after induction. There are 372 single cells in total. Normalized data was downloaded from GEO under accession number GSE52529 (file: GSE52529-fpkm-matrix.txt).
- *Treutlein et al. dataset:* This dataset was derived from Ref. 34. The experiment took samples from the developing mouse lung epithelium at embryonic days E14.5 (*n* = 45), E16.5 (*n* = 27), E18.5 (*n* = 83) and adulthood (*n* = 46), totalling 201 single cells. Normalized data was downloaded from GEO under accession number GSE52583 (file: GSE52583.Rda).
- *SemrauRB and SemrauSS datasets:* The two datasets named SemrauRB and SemrauSS were derived from Ref. 35, a study of retinoic acid driven differentiation of pluripotent mESCs to lineage commitment. SemrauRB used a recently developed Single Cell RNA Barcoding and Sequencing method (SCRB-seq), and consists of 425 single cell samples taken at 9 time points during mouse embryonic differentiation. There were 53 single cells taken at 0h. After all-trans retinoic acid (RA) exposure, 73 samples were taken at 6h, 49 samples at 12h, 27 samples at 24h, 72 samples at 36h, 43 samples at 48h, 54 samples at 60h, 29 samples at 72h and 25 samples at 96h. To increase cell numbers, 0-12h were considered to be pluripotent. SemrauSS uses a different scRNA-Seq technology (SMART-seq). A total of 292 single cells were taken at four time points (75 samples at 0h, 75 samples at 12h, 81 samples at 24h and 61 samples at 48h). Normalized data was downloaded from GEO under accession number GSE79578 (files: GSM2098545-scrbseq-2i.txt, GSM2098550-scrbseq-48h.txt, GSM2098555-smartseq-12h.txt, GSM2098546-scrbseq-6h.txt, GSM2098551-scrbseq-60h.txt, GSM2098556-smartseq-24h.txt, GSM2098547-scrbseq-12h.txt, GSM2098552-scrbseq-72h.txt, GSM2098557-smartseq-48h.txt, GSM2098548-scrbseq-24h.txt, GSM2098553-scrbseq-96h.txt, GSM2098549-scrbseq-36h.txt, GSM2098554-smartseq-2i.txt).
- *Wang et al. dataset:* This dataset was derived from Ref. 36, a study of non-directed differentiation over a 30-day period of neural progenitor cells (NPCs) into developing neurons, with the NPCs derived from hESCs. After quality control, there were 483 usable single cell samples describing the differentiation from neural progenitor cells (NPCs at day 0) to neurons and other differentiated cell-types. Cell numbers were 80 at day 0, 78 at day 1, 85 at day 5, 80 at day 7, 79 at day 10 and 81 at day 30. Normalized data was downloaded from GEO under accession number GSE102066 (file: GSE102066-normalized-counts.experiment1.DataMatrix.txt).
- *Yang et al. dataset:* This scRNA-Seq dataset was derived from Ref. 37, a study of differentiation of mouse hepatoblasts into hepatocytes and cholangiocytes. After quality control, 447 single-cell samples taken during embryonic development. There are 54 single-cells at embryonic day 10.5 (E10.5), 70 at E11.5, 41 at E12.5, 65 at E13.5, 70 at 14.5, 77 at 15.5 and 70 at E17.5. Normalized data was downloaded from GEO under accession number GSE90047 (file: GSE90047-Single-cell-RNA-seq-TPM.txt).
- *Yao et al. datasets 1-2:* This scRNA-Seq dataset (Yao1)was obtained by using a method based on multiplexed single-cell RNA-seq (CelSeq) and unique molecular identifiers (UMI)^14^. Cel-Seq was used to profile cells at multiple time points (days 0, 12, 19, 26, 40, and 54) in a study of differentiation of hESCs into cortical and non-cortical progenitor and finally into neurons. After quality control, there were 2684 single-cells: 40 hESCs, 504 collected at day-12, 278 at day-19, 595 at day-26, 502 at day-40 and 765 at day-54. Progenitor expansion into neurons occured at day-26. Normalized data was downloaded from GEO under accession number GSE86977 (file: GSE86977-UMI-20K.2684.csv). In addition, we also analysed a matched bulk RNA-Seq dataset (Yao2) from the same study, encompassing a total of 33 samples: 5 hESC samples (day-0), 3 samples at day-6, 4 samples at day-9, 6 samples at day-12, 5 samples at day-19, 6 samples at day-26 and 4 samples at day-54. Normalized data was downloaded from GEO under accession number GSE86985 (file: GSE86985-trueseq.tpm.csv).

### Further processing and quality control analysis

We used the following general procedure to further assess quality and further normalize the provided scRNA-Seq data. First, in all cases, we computed the coverage per single-cell, i.e. the number of genes with normalized counts (e.g. TPM/FPKM) above zero. Any single-cell with less than 10% coverage was removed. Next, we quantile normalized (QN) the resulting dataset to remove potential batch effects. For most datasets, we then renormalized the minimum read count to 1, i.e. any normalized count less than 1 was renormalized to 1. We justify this on the basis that the normalized counts of 1 and 0 are effectively indistinguishable. The threshold of 1 was derived by inspection of the distribution of normalized counts. After QN, we log2-transformed the data with an offset value of 0.1 added before log-transformation. The offset was added in order to ensure that the minimum value after log-transformation would not be 0, but a non-zero value (typically log_2_ 1.1 ≈ 0.13). We did not want zeroes in our final matrix to avoid singularities in the computation of the signalling entropy rate used in SCENT. The log-transformation was done to stabilize the variance and regularize the effects of outliers. Next, in the case of human data, we mapped gene symbols to Entrez gene IDs (if not annotated to Entrez gene IDs already), and finally averaged any entry mapping to the same Entrez gene ID. In the case of mouse, we first obtained the human homologs using the *annotationTools* Bioconductor package (http://www.bioconductor.org) and finally mapped to human Entrez gene IDs. The typical range of a resulting data matrix was then typically 0.13 to approx. 16.

We note that the Yao1 scRNA-Seq dataset used CelSeq-technology and UMIs, and was normalized using a different procedure as described in Ref. 14. For this dataset, we therefore used a slightly different choice of thresholds. First, the data downloaded was already on a log-scale, but still with a few significant outliers: thus, we regularized the data further by setting any values larger than 16 to 16. We then performed QN and finally any value less than 1 was set to a value of 0.1 (in order to remove zeroes from the data matrix).

The above procedure resulted in data matrices defined over human Entrez gene IDs and single-cells of following dimensions: Chu1 (18935 genes, 1018 single-cells), Chu2 (18935 genes, 758 single-cells), Chu3 (18935 genes, 19 bulk samples), Chu4 (18935 genes, 15 bulk samples), Trapnell (22351 genes, 372 single cells), Treutlein (12413 genes, 201 single-cells), SemrauRB (13755 genes, 425 single-cells), SemrauSS (13663 genes, 292 single-cells), WangNPC (8242 genes, 483 single-cells), YangML (16119 genes, 447 single-cells), Yao1 (22113 genes, 2684 single-cells), Yao2 (19659 genes, 33 bulk samples).

### Evaluation framework

In order to compare the different potency measures (described in the next subsections), we assessed their ability to discriminate single cells (and bulk samples) that ought to differ substantially in terms of their potencies. We reasoned that potency measures which would fail to discriminate such cell-types would be less preferable than those that are more robust. For each of the studies, we therefore devised comparisons between cell-types (or bulk samples) where typically we would expect to see a big difference in potency, so that it is reasonable to assume that higher AUC (discriminatory ability) and more significant P-values (as derived using one-tailed Wilcoxon rank sum tests) would indicate better potency measures. For the scRNA-Seq datasets, we devised a total of 11 independent comparisons, where the specific cell-types being compared and the corresponding cell numbers are given in **Table S1 in SI**. Below we briefly summarise the comparisons.

In Chu1, we chose to compare the pluripotent hESCs to all progenitor and non-pluripotent cell-types. In Chu2, we compared the hESCs at time 0 to the final timepoint (96h) corresponding to definite mesoderm/endoderm (i.e. non pluripotent). For the corresponding bulk sets Chu3 and Chu4, we chose the same groups, although for Chu4 only 12h and 24h were available to represent the pluripotent state, so for Chu4 we compared 12h/24h to 72h/96h, which yielded 6 samples in each group. Only using 12h and 96h would result in too few samples. We also note that the transition to the definite mesoendoderm occurs at 72h and beyond. In Trapnell, we compared the myoblasts at 0h to the differentiated skeletal muscle cells at 72h. In Treutlein, we compared embryonic day E14 to the adult differentiated cells. In SemrauRB, we compared cells at 0 and 12h to those harvested at 96h. We combined cells at 0h and 12h due to the treatment effect at 12h, not making 0h cells directly comparable to the rest. In SemrauSS, we compared cells at 0h to 48h. In WangNPC, we compared neural progenitors at 0 and 1 days, to differentiated neural cells at 30 days. We deemed safe to bunch together 0 and 1 day cells as differentiation occured much later in the assay and to increase cell numbers for the more potent NPC phenotype. In YangML, we compared embryonic day 10 (E10) to E17 single-cells. In Yao1, we devised 3 separate comparisons, because in this study there were many single-cells and because the nature of the experiment allowed precise definition of 3 potency states: hESCs at day-0, cortical and non-cortical progenitors at day-26 and differentiated neurons at 54-days. Thus, in Yao1a we compared day-0 to day-26, in Yao1b, we compared day-0 to day-54, and in Yao1c we compared day-26 to day-54. For the corresponding bulk sample study, Yao2, we also performed 3 comparisons, but in order to increase sample numbers we considered the pluripotent state to also includ day-9 samples, which is justified based on the findings of the original publication^14^.

### Adjusting for cell-cycle phase

Differences in cell potency between groups could be confounded by differences in the proportion of cycling vs non-cycling cells. For each single-cell we thus computed two cell-cycle scores (one for the G1-S and the other for the G2-M phase), using the procedure described in Ref. 8,38. Because the relation between potency and cell-cycle scores could be highly non-linear (as described in Ref. 8), linear adjustment procedures are inappropriate. Another approach to compare potency of cell-types, adjusted for cell-cycle, would be to set a threshold on the scores and only use cells with cell cycle scores below a given threshold. However, these thresholds may be study specific, and we found that if we were to use the same threshold for all studies, that this may lead for specific studies to highly unequal numbers of low-cycling cells in each comparison group, or to effectively no cells in a given group. For instance, in some studies we find that all single hESCs exhibit relatively high cycling scores. This indicates that disentangling cell-cycle effects and potency is extremely difficult, if not impossible, and may also be strongly dependent on the experimental details (e.g. culture conditions of cells etc.). Thus, we decided on a different approach which would allow us to compare cell-types, whilst adjusting for cell-cycle phase as much as possible. Specifically, for each study and cell-type of interest, we chose a lower quantile of cell-cycle scores based on the scores of all cells within that cell-type. Doing so, ensures that we are comparing sufficient numbers of single cells within each group, but also that we are using the lowest-cycling cells within each group. The quantiles were chosen to ensure sufficent numbers of cells in each comparison group. We note that since there are two cell-cycle scores, that we performed this operation on G1-S and G2-M separately, subsequently only using the single-cells that were in the lower defined quantile for both phases. Thus, absolute cell numbers could be low for some studies, but with the chosen quantiles enough cells were being compared to achieve potential statistical significance. The quantiles chosen, the number of low-cycling single-cells in each comparison group and the results obtained are listed in Table S3 in SI.

### Markov-Chain Entropy

We here describe the construction of Markov-Chain Entropy to represent the potency of a sample with a scRNA-Seq or bulk RNA-Seq profile. The construction requires two inputs, i.e. the normalized RNA-Seq profile and a fully connected signalling interaction network between the genes defined in the profile. For the network, we use the protein-protein interaction network. Specifically, we use the one with 8434 genes as used in Ref. 8. Given a network/graph *G* = {*V, E*} with *nV* nodes in *V* and *nE* edges in *E*, we model signaling as a particle on the network and let it do discrete time Markov transition. Denote the transition probability matrix as *P* = {*p*_*ij*_}, where *i, j*= 1, …, *n*_*E*_. We set *p*_*ij*_ = 0 if there is no edge between two different nodes *i* and *j*. ***p***_*ii*_ **can be nonzero, and is the probability of the particle staying at node *i***. We assume the network has a steady state with invariant distribution ***π*** = (*π*_1_, …, *π_nv_*) after the particle walks for long enough time. As we shall see below, the invariant measure or steady-state probabilities will be determined from the sample’s gene expression profile.

The *Markov-Chain Entropy* (MCE) of the Markov transition on network *G* is defined as

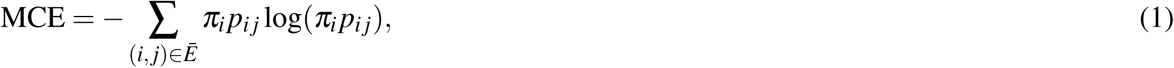

where the summation is only for the terms *p*_*ij*_ > 0, and *Ē* is the union set of *E* and all (*i, i*) self-loop edges. MCE can be split into two terms as

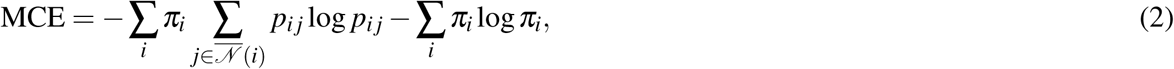

where 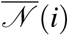 contains all the neighbors of node *i* including *i* itself.

The first term in Eq. (2) is named as the *edge entropy* (similar to but not the same as the average local entropy/signalling entropy in Ref. 8,39), and the second term is called the *node entropy*. Since *π*_*i*_*p*_*ij*_ means the probability (or information flow or actual interaction) transmitted from *i* to *j* along the directed edge (*i, j*) in *E*, MCE is indeed the *whole entropy* of the information flow induced by the Markov chain on the graph or network *G*, rather than the *entropy rate*^8^,^39^). Thus it can be thought as a measurement of the heterogeneity of a network in terms of actual interactions. The more homogeneous of interactions the graph is, the bigger its MCE grows. We think that this intuition can be utilized to characterize the stemness of cells.

### Computing MCE from single-cell RNA-seq dataset

Denote the dataset as *X* = {***x***(^1^),***x***(^2^), …,***x***(^*n*^)}, where *n* is the sample size and ***x***(^*i*^) is a length-nV column vector for gene expressions. *n*_*V*_ is not only the number of genes in consideration but also the counts of nodes in the gene network.

#### Choosing gene network

To compute the MCE, we must use the gene regulatory network (GRN) as the background. There exist many methods to estimate the GRN from gene expression dataset, like Graphical Lasso^40^,^41^, iterative clean-up calculation^42,43^, information theoretic approachescalculation^44^ and so on. However, generally, a large number of samples are needed to infer the direct edges accurately. The usual scRNA-seq datasets contain only from tens to hundreds samples, which are extremely few compared to thousands of genes. Thus in this work, we use the PPI network instead of gene regulatory network. The maximum connected component between PPI and the genes in dataset is denoted as A, which is the adjacency matrix used in the next steps.

#### Computing MCE for each sample

As mentioned above, we denote the 0-1 adjacency matrix of the PPI network as A, whose diagonal elements are also set to be 1s. For convenience, we still use *X* as the data matrix, and the size of *A* is *n*_*V*_ × *n*_*V*_. We have two assumptions before calculating the MCE for a single-cell sample ***x***:

1. The normalized gene expression 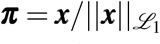 is the invariant distribution of a discrete time Markov transition matrix *P* on the gene network *A*;
2. Among all the possible transition matrix *P* with invariant distribution ***π*** and topology *A*, we choose the one with the largest entropy.

That means the MCE problem can be formulated as

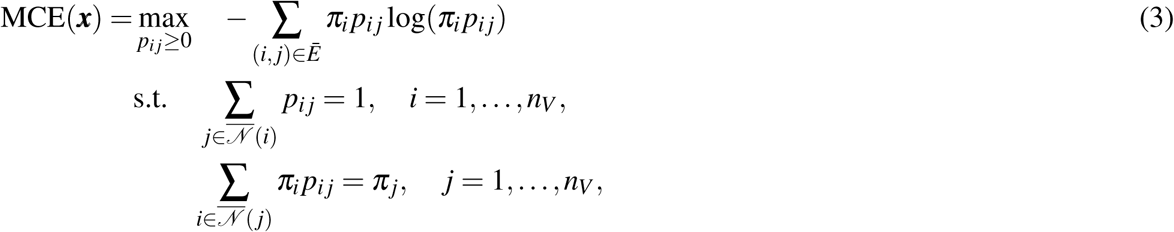

where the summation is only for the elements (*i, j*) ∈ *Ē* satisfying *A*_*ij*_ = 1, and 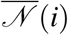 includes node *i* and all its neighbors.

Because ***π*** is known (so is the node entropy), the above can be written into a standard convex optimization problem for edge entropy:

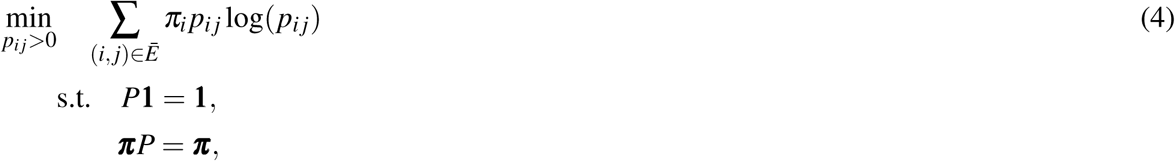

where *P* = (*p*_*ij*_)*n*_*V*_ × *n*_*V*_ has the same topology as A and variables only are located at the position of 1s in A. Though the problem Eq. (4) is convex with linear constraints, there are about 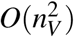 variables, which means to find an almost million-dimension optimizer for one cell! It sounds like infeasible for the first look, but luckily we can change the variables into its Lagrangian multipliers and solve it practically like:

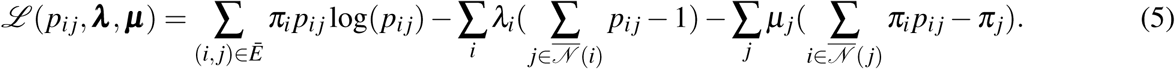

Calculate the first derivations of (5) we get

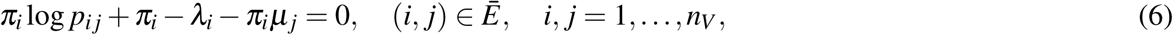

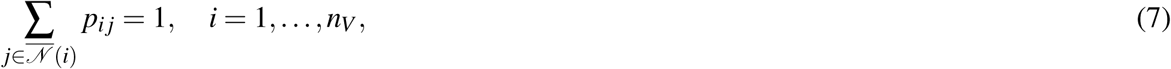

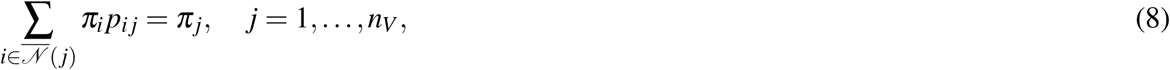

where only the terms *p*_*ij*_ ≥ 0 corresponding to *A*_*ij*_ = 1 are used in the equations. Denote *α*_*i*_ = exp(*λ*_*i*_/*π*_*i*_), *β*_*j*_ = exp(*μ*_*j*_ — 1) and *p*_*ij*_ = *α*_*i*_*β*_*j*_*a*_*ij*_ then we have transformed the optimization problem into solving nonlinear equations.

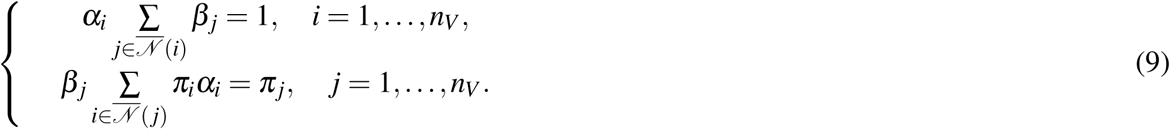

There are fortunately *2nV* nonlinear equations corresponding to 2*n*_*V*_ variables.

We propose an iteration method to solve the equations (9) as

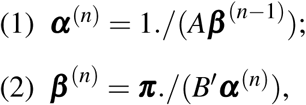

where *A* is the adjacency matrix, *B* is got by multiplying each element in row *i* of *A* by ***π***_*i*_ (i.e. *b*_*ij*_ = ***π***_*i*_), and .“/” means division by elements. With a proper initial point ***β***^(0)^, we can do the iteration until (***α***^(*n*)^, ***β***^(*n*)^) converging into a fixed point (***α***^*^, ***β***^*^) Then the MCE of a single-cell sample can be calculated as

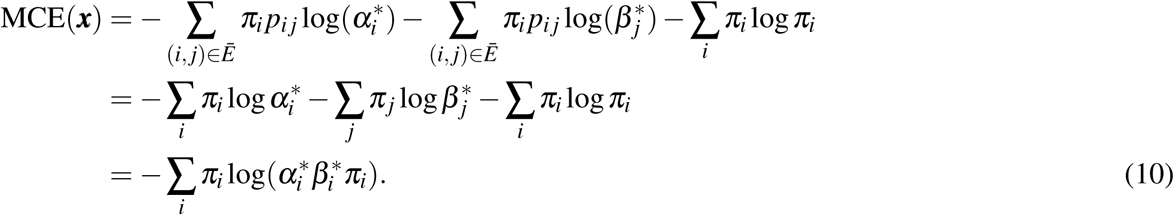

#### Normalization of MCE

To normalize the MCE into [0,1], we should compute the maximum possible MCE of all the networks with topology A. According to the formation of MCE in Eq. (1), the maximum is reached when *π*_i_ ~ *d*_*i*_ and *p*_*ij*_ = 1/*d*_*i*_, where *d*_*i*_ is the degree of node *i* (including *i* itself). Denote *d* = Σ_*i*_ which is equal to the count of nonzero terms in *A*. Thus we have

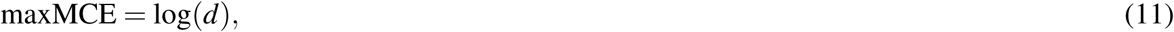

and the normalized MCE is computed as

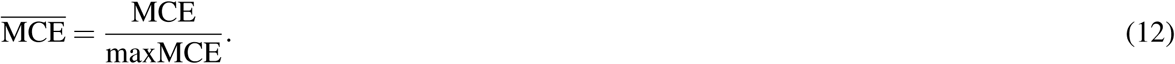

## Other entropy indices

### Signalling entropy

Signalling entropy (SR) was proposed in Ref. 5,8,39. When computing SR, the PPI network should be provided too, while the weights on edges are supposed to be proportional to the normalized expression levels of the genes, which means *w*_*ij*_ ~ *x*_*i*_*x*_*j*_. Transition matrix *P* has entries

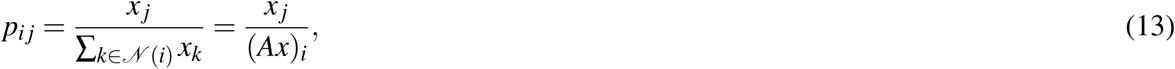

where *N*(*i*) means the neighbors of node *i* and *A* is the PPI adjacency matrix. With detailed balance condition and invariant measure ***π*** computed from *P* as 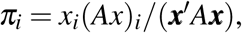 signalling entropy is defined and calculated as

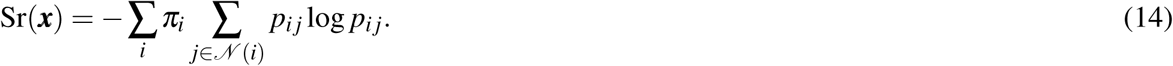

To normalize the signalling entropy, maximun entropy is computed by

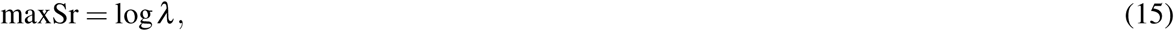

where *λ* is the largest eigenvalue of adjacency matrix *A*. Normalized entropy rate is defined as

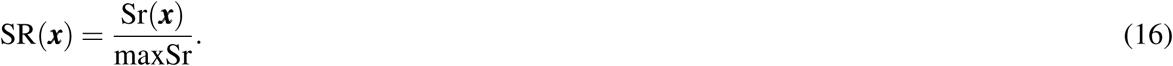

The main difference between SR and MCE are:

1. The adjacent matrix *A* has zero diagonal elements in SR, while it has ones in MCE. The basis of SR is interaction between proteins through the law of mass action, while the basis of MCE is a Markov transition on the gene regulatory network. The particle in MCE has the probability to stay in one node which is much meaningful.
2. (2) For MCE, we optimize the entropy to get the most possible transition matrix with fixed invariant measure. Maximum entropy means the most possible case, while the weight of stochastic matrix *P* in SR is computed by law of mass action which is a determined method.

### StemID

Grün *et al.* proposed the StemID algorithm to estimate differentiation potency of single cells in Ref. 6. It does not use a PPI network, but has to cluster the cells into groups. StemID computes a score *s* for each cluster as

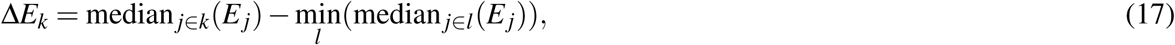

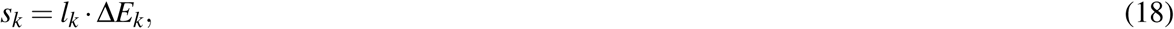

where *k* labels a cluster, *l*_*k*_ is the number of significant links, *E*_*j*_ is the entropy of cell *j* defined as 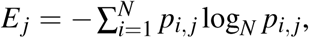 *N* is the total number of genes and *p*_*i, j*_ is the normalized number of reads mapping to gene *i* in cell.

To compare fairly and objectively to other entropy indices like SR and MCE, we take *E*_*j*_ as the potency estimate from StemID for each cell, which is much like the node entropy defined in Eq. (2) and just like the comparison done in Ref. 8. Importantly, we note that the computation of StemID was performed on the normalized (unlogged) counts, after quantile normalization. This is line with how it was recommended to compute StemID in the original paper^6^.

### SLICE

SLICE was another entropy based index proposed in Ref. 7. It first clusters genes into *m* functional groups according to GO-terms. Then for each cell *j*, SLICE computes the entropy of the active distribution of the groups as 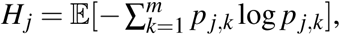 where *p* _*j, k*_ is the activity score of group *k* in cell *j* and the expectation is taken over a number of bootstraps. We computed SLICE as described in Ref. 8 using the R-functions provided by Guo et al.^7^. We note that, as with StemID, the computation of SLICE entropy is supposed to be done on the unlogged normalized counts. Thus, we used the downloaded normalized count data matrices and only performed quantile normalization before estimating SLICE entropy. The expression threshold parameter in SLICE was set to 1 (default value) for all studies, expect for Yao1 where the data was already regularized (logged) and where we used a threshold of 0.1. We note that in the Yao1 study data had been renormalized by subsampling single-cells to 20,000 UMIs^14^.

### Construction of the landscape

We construct an illustrative landscape in Fig. 1B of the main text.

Firstly, *X* and *Y* axes are the first two coordinates computed by t-SNE.^45^ t-SNE is a dimension reduction tool used widely in many subjects now. It can make different clusters of cells more separated and thus more clear to be observed in low dimensions. The z-axis is MCE value. In the tSNE-MCE figure, higher cells are more pluripotent as they have higher entropy.

To plot an illustrative potential landscape, we use a function like

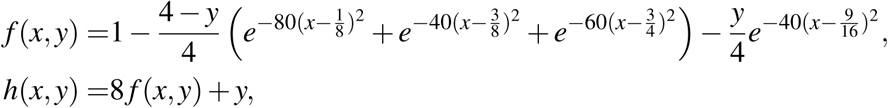

and plot the height values of *h(x, y)* for *(x, y)* ∈ [0,1] × [-1,4]. The idea of constructing such landscape function is to make it a Gaussian mixture in *x*-direction but linear in *y*-direction.

At last, we put the surface at the right place in the tSNE-MCE figure. We have to mention that the landscape is an illustration like Waddington’s in order to show how cells differentiate from stem cells to others although their MCEs are the actual values based on our method from the data. Our purpose is to show high MCE indeed presents the cell–s pluripotent ability.

